# Cell Type-Specific and Diabetic Kidney Disease-Associated Expression of Long Non-Coding RNAs in Human Kidneys

**DOI:** 10.1101/2025.01.14.632910

**Authors:** Juliette A de Klerk, Roderick C Slieker, Wilson C Parker, Haojia Wu, Yoshiharu Muto, Rudmer J Postma, Leen M ’t Hart, Janneke HD Peerlings, Floris Herrewijnen, Heein Song, H Siebe Spijker, Sébastien J Dumas, Marije Koning, Loïs AK van der Pluijm, Hans J Baelde, Tessa Gerrits, Joris I Rotmans, Anton Jan van Zonneveld, Coen van Solingen, Benjamin D Humphreys, R Bijkerk

## Abstract

**Background:** Long non-coding RNAs (lncRNAs) play essential roles in cellular processes, often exhibiting cell type-specific expression and influencing kidney function. While single-cell RNA sequencing (scRNA-seq) has advanced our understanding of cellular specificity, past studies focus solely on protein-coding genes. We hypothesize that lncRNAs, due to their cell-specific nature, have crucial functions within particular renal cells and thereby play essential roles in renal cell function and disease.

**Methods:** Using single-nucleus RNA sequencing (snRNA-seq) data from kidney samples of five healthy individuals and six DKD patients, we explored the non-coding transcriptome. Cell type-specific lncRNAs were identified, and their differential expression in DKD was assessed. Integrative analyses included expression quantitative trait loci (eQTL), genome-wide association studies (GWAS) for estimated glomerular filtration rate (eGFR), and gene regulatory networks. Functional studies focused on *TARID*, a lncRNA with podocyte-specific expression, to elucidate its role in podocyte health.

**Results:** we identified 349 lncRNAs with cell type-specific expression across kidney cell types. Of these, 104 lncRNAs were differentially expressed in DKD. Integrative analyses, including eQTL data, GWAS results for eGFR and gene regulatory networks, pinpointed *TARID*, a podocyte-specific lncRNA, as a key candidate upregulated in DKD. Functional studies confirmed *TARID*’s podocyte-specific expression and revealed its central role in actin cytoskeleton reorganization, a critical process in podocyte health.

**Conclusions:** Our study provides a comprehensive resource of single-cell lncRNA expression in the human kidney and highlights the importance of cell type-specific lncRNAs in kidney function and disease. Specifically, we demonstrate the functional relevance of *TARID* in podocyte health. This work underscores the utility of integrating scRNA-seq with functional genomics to uncover novel regulatory mechanisms in kidney biology.

**Key points:**

- This provides a resource for kidney (cell type-specific) lncRNA expression and demonstrates the importance of lncRNAs in renal health.
- We identified 349 cell type-specific lncRNAs in the human kidney, with 104 showing altered expression in diabetic kidney disease (DKD).
- *TARID*, a podocyte-specific lncRNA upregulated in DKD, is crucial for actin cytoskeleton reorganization in podocytes.

## Introduction

Kidney disease remains a major health challenge with significant morbidity and mortality worldwide [1]. Genome-wide association studies (GWAS) have provided valuable insights into the genetic underpinnings of kidney disease, highlighting key loci and potential biomarkers for disease susceptibility [2]. Recent research has further expanded our understanding of the human genome, revealing that approximately 76 – 97% of it consists of non-coding sequences [3]. Among these, long non-coding RNAs (lncRNAs) have emerged as key regulatory molecules involved in a wide range of cellular processes, including gene expression, epigenetic modulation, and disease pathogenesis [4]. Notably, lncRNAs are increasingly recognized for their cell type-specific functions [5]. Numerous lncRNAs have been implicated in kidney function regulation. Among them, Metastasis Associated Lung Adenocarcinoma Transcript 1 (*MALAT1*) stands out as a significant regulator in diabetic kidney disease (DKD) [6]. Silencing *MALAT1* with small interfering (si)RNA has been shown to partially restore podocyte function, highlighting its role in kidney health [7]. Similarly, Nuclear paraspeckle Assembly Transcript 1 (*NEAT1*) is associated with renal inflammation and fibrosis, especially in DKD and chronic kidney disease (CKD), where it exacerbates inflammation by modulating immune-related gene expression [8]. Additionally, the kidney-specific TGF-β/SMAD3-interacting long non-coding RNA (*lnc-TSI*), inhibits renal fibrogenesis by impeding the TGF-β/SMAD3 pathway [9]. Despite the growing interest in kidney-related lncRNAs, the specific functions and mechanisms of many remains poorly understood, in part due to limited knowledge of their kidney cell type-specific expression and complex regulatory roles. Moreover, only a fraction of kidney-associated lncRNAs has been studied, leaving much of the lncRNA landscape in renal biology unexplored.

Single-cell RNA sequencing (scRNA-seq) as well as single nucleus RNA-sequencing (snRNA-seq) offer unprecedented resolution for exploring the transcriptomic landscape of the kidney and other tissues, enabling the dissection of gene expression profiles at the level of individual cell types. While previous studies have primarily focused on protein-coding genes, the rich lncRNA content of these datasets remains underexplored [10]. Given the tissue specificity and functional significance of lncRNAs in kidney physiology, a focus on lncRNAs in scRNA-seq datasets is warranted to uncover their roles in both normal kidney function and disease states, including DKD.

This study aims to reassess a snRNA-seq dataset of the human kidney [10], placing particular emphasis on the identification and characterization of lncRNAs. By leveraging the high-resolution data from snRNA-seq, we seek to determine cell type-specific expression of lncRNAs, investigate their potential associations with DKD, and integrate this information with genetic and regulatory network data to identify key lncRNAs involved in kidney function and pathology. Specifically, we focus on cell type-specific lncRNAs associated with DKD, a major complication of diabetes, which leads to progressive kidney dysfunction and is a leading cause of end-stage kidney disease worldwide [11]. Through this reanalysis, we aim to shed light on the underexplored landscape of lncRNAs in the kidney, providing novel insights into their potential roles in disease progression and offering new avenues for therapeutic targeting in DKD and other kidney-related disorders.

## Methods

Detailed methods are available in the **Supplemental Materials**. In short, this study reanalysed our snRNA-seq dataset from human kidney cortex samples of five control and six DKD patients to investigate lncRNAs. Key analyses included cell-type-specific and DKD-associated lncRNAs identified using Seurat, gene regulatory networks via Pearson correlations, and pathway enrichment with Reactome and KEGG. *TARID* expression was further examined using GWAS-eQTL analyses, *in situ* hybridization, kidney organoid scRNA-seq datasets, qPCR on glomeruli samples, and in vitro experiments in cultured podocytes with *TARID* knockdown. Functional effects of *TARID* knockdown on podocyte morphology and gene expression were evaluated through imaging, bulk RNA-seq, and pathway analysis. Statistical analyses confirmed significant findings, highlighting *TARID*’s potential role in kidney function and disease.

## Results

### Single-nucleus RNA-sequencing identifies cell type-specific lncRNAs

SnRNA-seq data of five controls and six DKD individuals were reanalysed. The snRNA-seq dataset included 39,176 cells, encompassing all key cell types present in the kidney cortex: proximal tubule, PT, VCAM1(+) proximal tubule, PT_VCAM1, parietal epithelial cells, PEC, ascending thin limb, ATL, CLDN16(-) thick ascending limb, TAL1, CLDN16(+) thick ascending limb, TAL2, early distal convoluted tubule, DCT1, late distal convoluted tubule, DCT2, principal cells, PC, type A intercalated cells, ICA, type B intercalated cells, ICB, podocytes, PODO, endothelial cells, ENDO, mesangial cells and vascular smooth muscle cells, MES, fibroblasts, FIB, leukocytes, LEUK (**Fig. 1A**). Original UMAP clustering was preserved [10]. A total of 17,910 lncRNAs are available in GENCODE version 32 (**ESM Table 1**), of which 10,327 lncRNAs were present in the snRNA-seq, encompassing 37% of the total amount of transcripts. The remaining transcripts mapped to other types of transcripts including protein coding. Most cell types have around 15% of the total transcripts mapped to lncRNAs (**Fig. 1B-C**). We found that 349 lncRNAs exhibited cell type enriched or specific expression in 15 cell types (**Fig. 1D, ESM Table 2**). Most cell type specific lncRNAs were specific for the PT cluster, followed by TAL1 and PODO. We observed a correlation between the number of cells in a cluster and the number of cell type specific lncRNAs in a cluster (R^2^ = 0.78, P-value = <0.001) (**ESM Fig. 1**). However, some cell types have more cell specific lncRNAs then expected based on the number of cells in that cluster, for example in podocytes. Fibroblasts were the only cell type without (statistically significant) cell type specific expression. The expression of individual cell type–specific lncRNAs indicated a high correlation with canonical protein-coding marker genes (**Fig. 1E**). For example, expression of *TARID*, which was a specifically expressed lncRNA in podocytes, displayed a strong correlation to the expression of *NPHS2* (podocin), a representative marker gene of podocytes. We identified 405 lncRNAs associated with DKD within a specific cell type. Of these, 190 lncRNAs were unique, with several exhibiting differential expression across multiple cell types. Notably, this set also includes X-inactive specific transcript (*XIST*), a well-characterized lncRNA involved in X-chromosome inactivation [12]. Of these DKD associated lncRNAs, 104 demonstrated cell-type-specific expression (**Fig. 2A, ESM Table 3-4**). We observed that the endothelial-specific lncRNA *PCAT19* is upregulated in DKD, consistent with its previously reported role in vascular injury [13]. Additionally, we identified several novel lncRNAs with specific expression patterns. For example, lncRNA *AC026333.3* is expressed exclusively in type A intercalated cells and is downregulated in DKD, while *AC003044.1* is uniquely expressed in both VCAM1-positive and -negative proximal tubule cells and is significantly upregulated in DKD. Furthermore, lncRNA *AL355612.1*, primarily expressed in early distal convoluted tubule cells and to a lesser extent in late distal convoluted tubule cells, is also downregulated in DKD. We found that lncRNA *AC092813.2* shows podocyte-specific expression and is upregulated in DKD. Another lncRNA, *NEAT1*, is expressed in all cell types but exhibits the highest expression in proximal tubule cells, with even greater expression in DKD (**ESM Fig. 2A-F**). Notably, *NEAT1* has been previously implicated in DKD pathogenesis [14]. Interestingly, certain cell types demonstrate more pronounced cell-specific expression of lncRNAs. For instance, we identified five lncRNAs exclusively expressed in podocytes, with no expression detected in other cell types: *TARID*, *AL024495.1*, *AC064875.1*, *SLC7A14-AS1* and *AC092813.2*. All of them are significantly upregulated in DKD (**Fig. 2A-B**).

**Figure 1.**
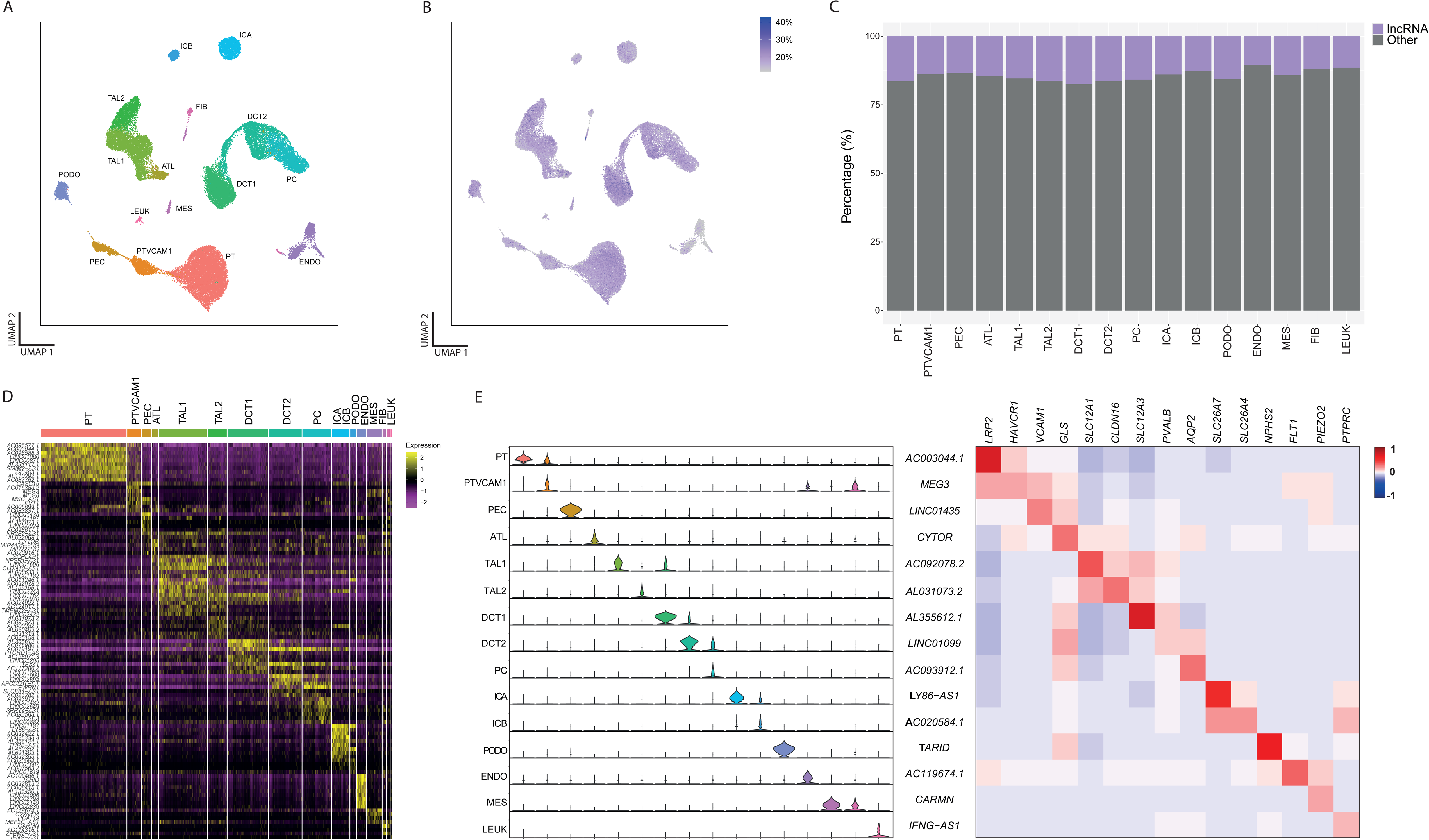
SnRNA-seq of human kidney reveals cell type-specific lncRNAs. (**A**) UMAP visualization of snRNA-seq data, highlighting overall cellular clustering. (**B**) UMAP projection illustrating the distribution of lncRNAs within the snRNA-seq dataset. (**C**) Distribution of lncRNAs across different cell types. (**D**) Identification of 349 cell type-specific lncRNAs, with the top five lncRNAs for each cell type displayed. (**E**) Analysis of cell type-specific lncRNAs and their correlation with known protein-coding marker genes. Left: Violin plots showing expression patterns of representative lncRNAs in specific cell types. Right: Heatmap depicting the correlation between these lncRNAs and cell type-specific protein-coding marker genes, with colour intensity indicating the degree of correlation.

**Figure 2.**
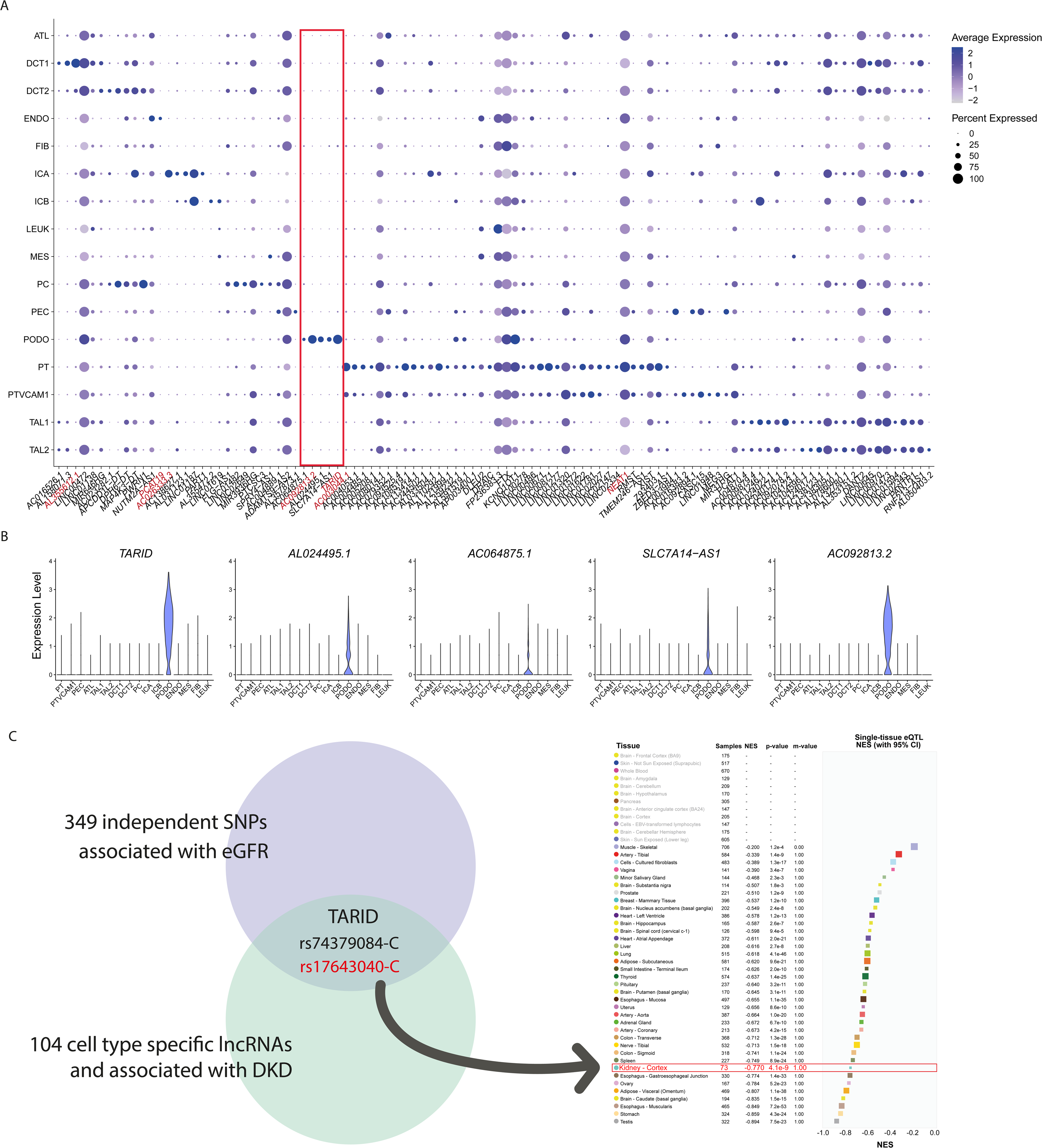
Cell type-specific and DKD-associated lncRNAs overlap with eGFR-associated loci. (**A**) Dot plot showing 104 lncRNAs that are both cell type-specific and associated with diabetic kidney disease (DKD). LncRNAs discussed in the main text are highlighted in red: *AL355612.1*, *PCAT19*, *AC026333.3*, *AC092813.2*, *TARID*, *AC003044.1* and *NEAT1.* Red box indicates podocyte-specific transcripts. (**B**) Violin plot of five podocyte-specific and DKD associated lncRNAs: *TARID*, *AL024495.1*, *AC064875.1*, *SLC7A14-AS1* and *AC092813.2*. (**C**) Among 349 independent SNPs significantly associated with eGFR, only one lncRNA, *TARID,* was located within the 1MB region. The SNPs rs74379084-C and rs17643040-C were found in this region, with rs17643040-C identified as a cis-eQTL in multiple tissues, including the kidney cortex, for *TARID*.

### eQTL/GWAS analysis identifies podocyte-specific lncRNA TARID

To identify lncRNAs in our dataset that may be causally related to kidney function, we next analysed a GWAS dataset that investigated single nucleotide polymorphisms SNPs in relation to kidney function. Among 349 independent SNPs that are significantly associated with estimated glomerular filtration rate (eGFR) (P-value < 5 x 10^−8^) from GWAS (GWAS Catalog no. GCST90019506), only one SNP fell in the 1MB centered region of the 104 lncRNAs which were cell type specific and associated with DKD: *TARID* (rs74379084-C, and rs17643040-C) (**ESM Table 5**). Of these 2 SNPs, rs17643040-C was a cis-eQTL in multiple tissues including the kidney cortex for *TARID*. This cis-eQTL is associated with lower *TARID* expression in the kidney cortex (P-value = 4.1 x 10^-9^, normalized effect size = -0.77) and higher eGFR (P-value = 3 x 10^-9^, beta = 0.0199, CI = 0.013 - 0.027) (**Fig. 2C**). Interestingly, *TARID* was exclusively expressed in almost all podocytes and not in other cell types (**Fig. 3A**) and significantly upregulated in DKD (adjusted P-value = 1.3 x 10^-9^) (**Fig. 3B**).

**Figure 3.**
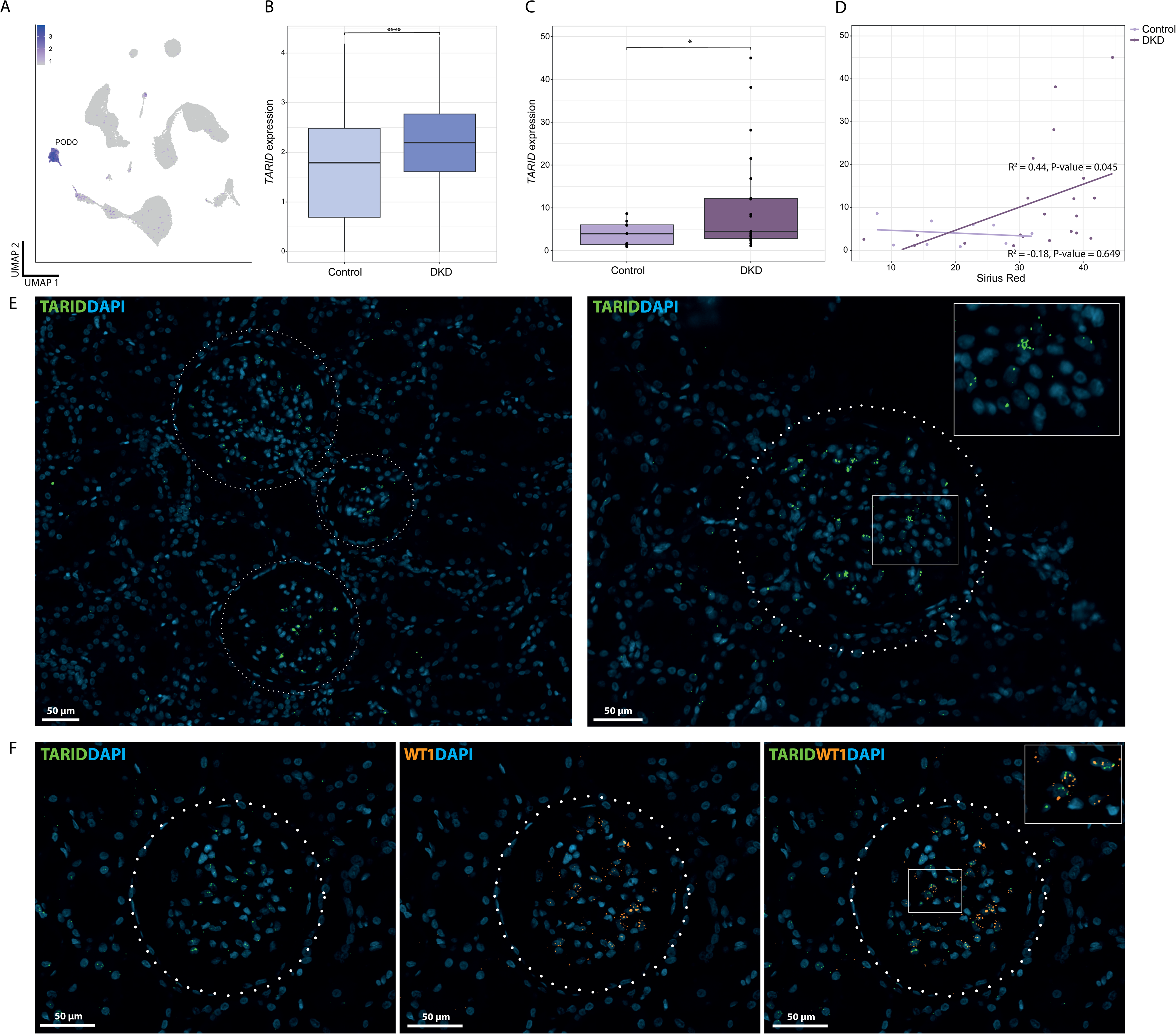
*TARID* is a podocyte-specific lncRNA. (**A**) UMAP projection showing TARID expression across kidney cell types. (**B-D**) Expression of TARID in healthy kidneys and in kidneys of diabetic kidney disease patients (DKD) analysed by snRNA-seq (**B**), by qPCR analysis of additional RNA samples (n = 32) from glomerular kidney biopsies from patients with DKD and healthy controls (**C**) and correlation with Sirius red (**D**). (**E**) *In situ* hybridization of *TARID* (green) in human FFPE kidney tissue. The glomeruli is displayed within the dotted line, bar□=□50□μm. Left and right display different glomeruli on different slides. Dapi (blue) was used to stain for nucleus. (**F**) *In situ* hybridization of *TARID* (green) and *WT1* (orange) and their colocalization in human FFPE kidney tissue. The glomeruli is displayed within the dotted line, bar□=□50□μm. Dapi (blue) was used to stain for nucleus.

### Confirmation of increased TARID levels in validation cohort of glomerular RNA samples

To confirm the upregulation of *TARID* in DKD, RNA samples from glomerular kidney biopsies of patients with DKD were analysed for *TARID* expression. We observed a significant upregulation of the lncRNA *TARID* in patients with DKD compared to control (P-value = 0.01), which is in line with the snRNA-seq analysis (**Fig. 3C**). Furthermore, *TARID* expression showed correlation with Sirius red staining in DKD samples (R^2^ = 0.44, P-value = 0.045) and not in control samples (R^2^ = -0.18, P-value = 0.65) (**Fig. 3D**). Sirius Red is a well-established marker of interstitial fibrosis and a crucial indicator of disease progression.

### Validation of Podocyte-Specific TARID Expression in Human Kidneys and Kidney Organoids

To validate the podocyte-specific expression of *TARID* indicated by snRNA-seq data, we performed RNAscope analysis. *TARID* showed specific staining in glomerular cells, that based on their location most likely represent podocytes, within human kidney sections (**Fig. 3E**). Additionally, co-staining with the podocyte marker Wilms Tumor 1 (*WT1*) demonstrated partially co-localization of *TARID* and *WT1* in glomerular podocytes, further confirming the podocyte-specific expression of *TARID* (**Fig. 3F**). Furthermore, in two independent scRNA-seq datasets of kidney organoids—comprising both untransplanted and transplanted kidney organoids, we assessed the specific expression of *TARID* [15]. In both datasets, *TARID* displayed podocyte-specific expression, which was consistently observed across all developmental stages of these cells (**ESM Fig. 3**). In addition, podocyte-specific expression of *TARID* was also confirmed in other scRNA-seq datasets of the human kidney/kidney organoid in the Kidney Interactive Transcriptome (KIT) database (**ESM Fig. 4, ESM Table 6**).

### Gene regulatory networks indicate TARID associates with podocyte morphology

We set up a gene regulatory network to explore the functional role of *TARID* in podocytes. This was done by correlating *TARID* with all other genes in the snRNA-seq specifically in podocytes using the Pearson correlation coefficient (**ESM Table 7**). Eight of the 12 most highly correlated genes with *TARID* expression (Pearson correlation coefficient ≥ 0.5): *PARD3B (***Fig. 4A**), *ST6GALNAC3*, *COL4A3*, *PLCE1*, *MAGI2*, *CCDC91*, *NEBL* and *IQGAP2* have been linked to podocyte dysfunction in literature (**Fig. 4B**). The genomic locus of *TCF21* Antisense RNA Inducing Promoter Demethylation (*TARID*) lies adjacent to the *TCF21* gene, transcribed in the opposite direction (**ESM Fig. 5**). Previous studies have demonstrated that TARID plays a critical role in activating TCF21 expression by promoting promoter demethylation, suggesting its involvement in regulating gene expression through epigenetic modifications [16]. We found that 101 genes had a Pearson correlation coefficient ≥ 0.4. However, it is notable that *TCF21* itself was not included in this correlation, despite its close proximity to *TARID*. Pathway enrichment with ReactomePA and KEGG on the correlated genes pointed out enrichment for: *Rho GTPase cycle* (**Fig. 4C, ESM Table 8**) and *Regulation of the actin cytoskeleton* (**Fig. 4D, ESM Table 9**). Additionally, using EnrichR we found enrichment for *Abnormal podocyte foot process morphology*, *Podocyte foot process effacement* (MGI Mammalian Phenotype Level 4 2024) and *Primary focal segmental glomerulosclerosis*, *Nephrotic syndrome* (WikiPathways 2024 Human) (**ESM Table 10**) [17–19].

**Figure 4.**
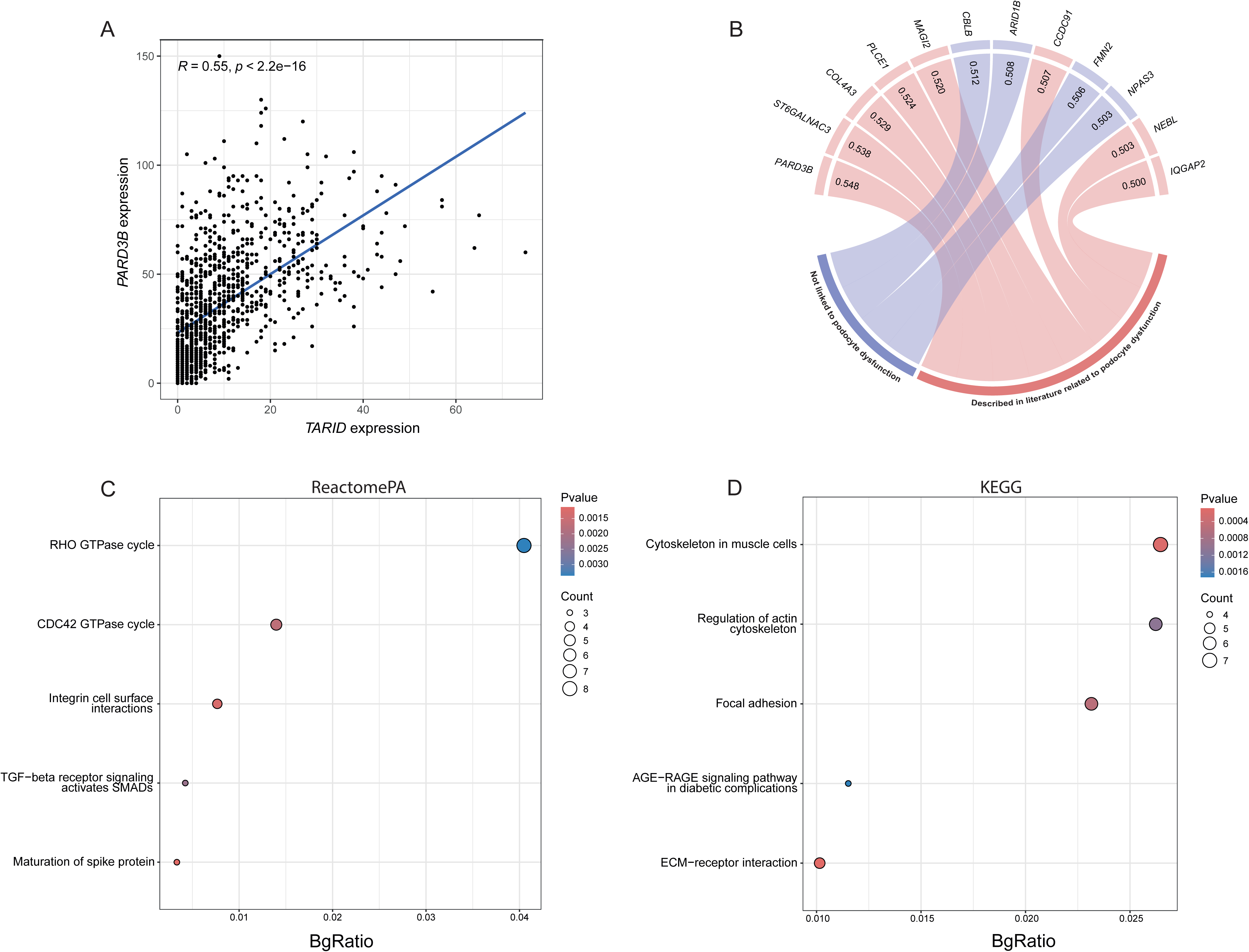
Gene regulatory network of *TARID*. (**A**) Correlation analysis of *PARD3B* and *TARID* in podocytes. (**B**) Circos plot showing correlation coefficient (> 0.5) for *TARID* with genes that are linked to podocyte dysfunction (red) and not previously linked to podocyte dysfunction (blue). (**C-D**) Pathway enrichment analysis (genes = 101, Pearson correlation coefficient > 0.4) with Reactome Pathway Analysis (PA) (**C**) or Kyoto Encyclopedia of Genes and Genomes (KEGG) (**D**). Colour of dots indicate P-value, and size of dots indicate gene count. X-axis shows BgRatio.

### Antisense LNA Gapmer knockdown of TARID in podocytes

Next, we sought to further explore the function of *TARID* in podocytes. To that end, *TARID* expression was downregulated with three different antisense LNA Gapmer designs in conditionally immortalized human podocytes (CIHP-1). One Gapmer seemed most effective in knockdown of *TARID* expression, Gapmer design 2 (**ESM Fig. 6**). Using this *TARID* Gapmer for further studies, next eight samples were selected for bulk RNA-sequencing (NC Gapmer n = 4, *TARID* Gapmer 2 n = 4). Both qPCR and bulk RNA-seq revealed efficient knockdown of *TARID* (**Fig. 5A, ESM Fig. 7**). Our results show that *TARID* knockdown leads to a modest reduction (16%, not significant) in *TCF21* gene expression (**ESM Fig. 8**). In total, we found 258 statistically significant differentially expressed genes upon *TARID* knockdown, including 171 downregulated and 87 upregulated genes (**Fig. 5B, ESM Table 11**), of which 44 are podocyte specific in our snRNA-seq dataset (**ESM Table 12**). Among the differentially expressed genes were, *NOTCH3* and *WNT2*, which are two well-known genes associated with kidney and podocyte function. Of the 101 genes that correlated with *TARID* in the snRNA-seq (Pearson correlation coefficient ≥ 0.4) we found 3 genes that also significantly associated with *TARID* knockdown with similar direction of effect, *SPOCK1*, *SULF1* and *NEDD9* (**ESM Fig. 9**). Of which, *SULF1* is associated with podocyte foot process effacement and *NEDD9* is an actin related gene [20, 21]. In line with the snRNA-seq-based *TARID* gene regulatory network analysis, pathway enrichment analysis on differentially expressed genes upon *TARID* knockdown revealed genes related to among others *cytoskeleton reorganization* (**Fig. 5C, ESM Table 13**). Further investigation of the genes associated with cytoskeleton reorganization revealed a specific involvement with the actin cytoskeleton (**ESM Table 14, ESM Fig. 10**). As such, the F-actin cytoskeleton organization was measured after knockdown of *TARID* in CIHP-1 cells. Indeed, we confirmed that knockdown of *TARID* results in altered actin cytoskeleton structure (**Fig. 6A**). We observed an upregulation in the total length of the actin filaments (P-value = 0.02, **Fig 6B**). Furthermore, we also observed a upregulation in the total number of actin filaments upon *TARID* knockdown (P-value = 0.03, **Fig 6C**).

**Figure 5.**
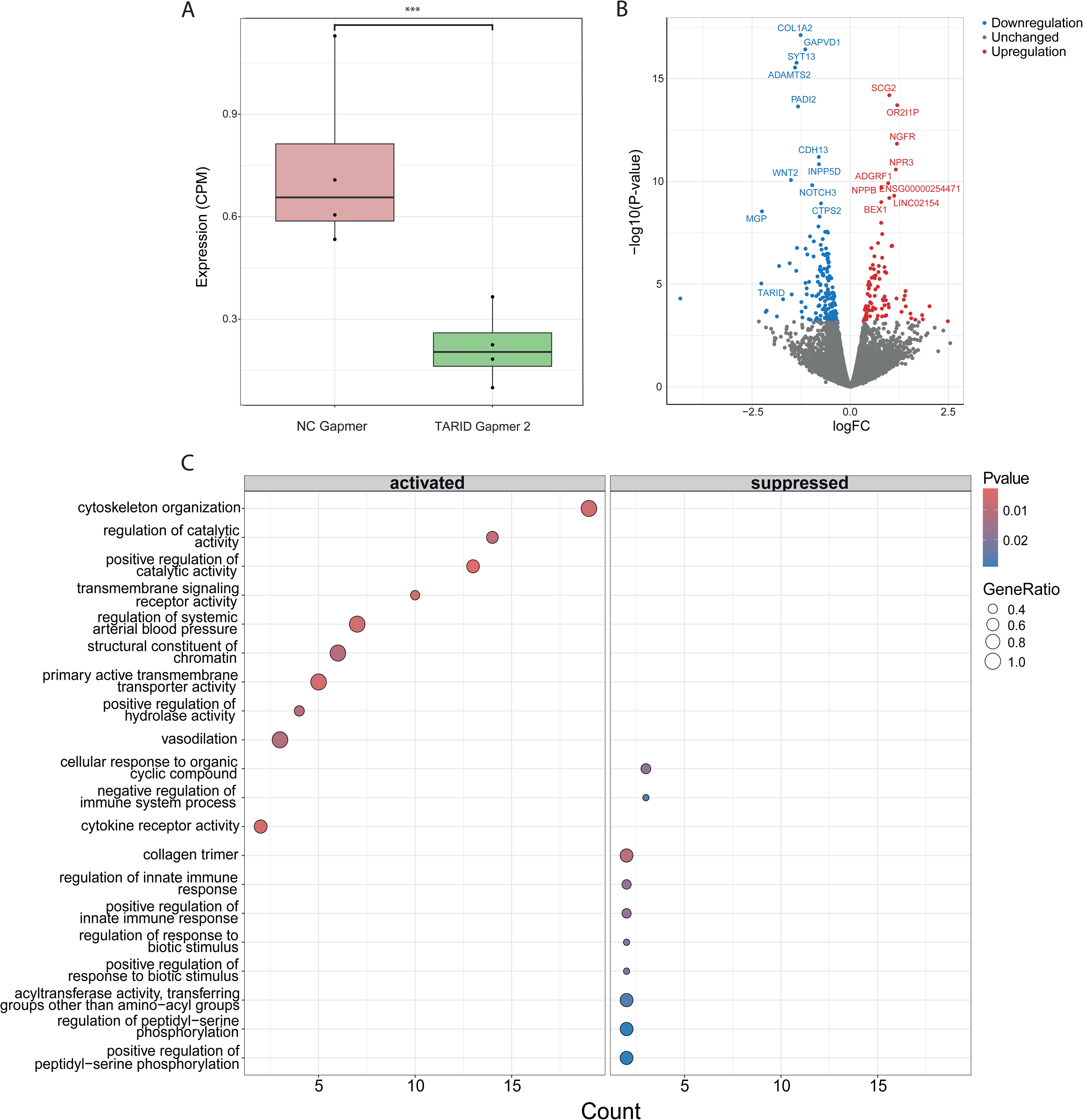
Bulk RNA-seq analysis of podocytes following *TARID* knockdown. (**A**) RNA-seq analysis of *TARID* expression in CIHP-1 cells upon transfection with scrambled Gapmer and *TARID* Gapmer (design 2). (**B**) Volcano plot displaying differentially expressed genes in CIHP-1 cells following *TARID* knockdown. Blue dots indicate down regulated genes (n = 171), red dots indicates upregulated genes (n = 87). Grey dots indicated not-significant changed genes. Gene names of the top 20 associated genes and *TARID* are displayed in the volcano plot. (**C**) Pathway enrichment analysis results of significantly differentially expressed genes. Left panel indicates activated pathways. Right panel indicates suppressed pathways. Colour of dots indicate P-value, and size of dots indicate GeneRatio. X-axis shows Count. (**D**) Microscopic images of CIHP-1 cells transfected with scrambled Gapmer or TARID Gapmer (design 2) stained for F-actin.

**Figure 6.**
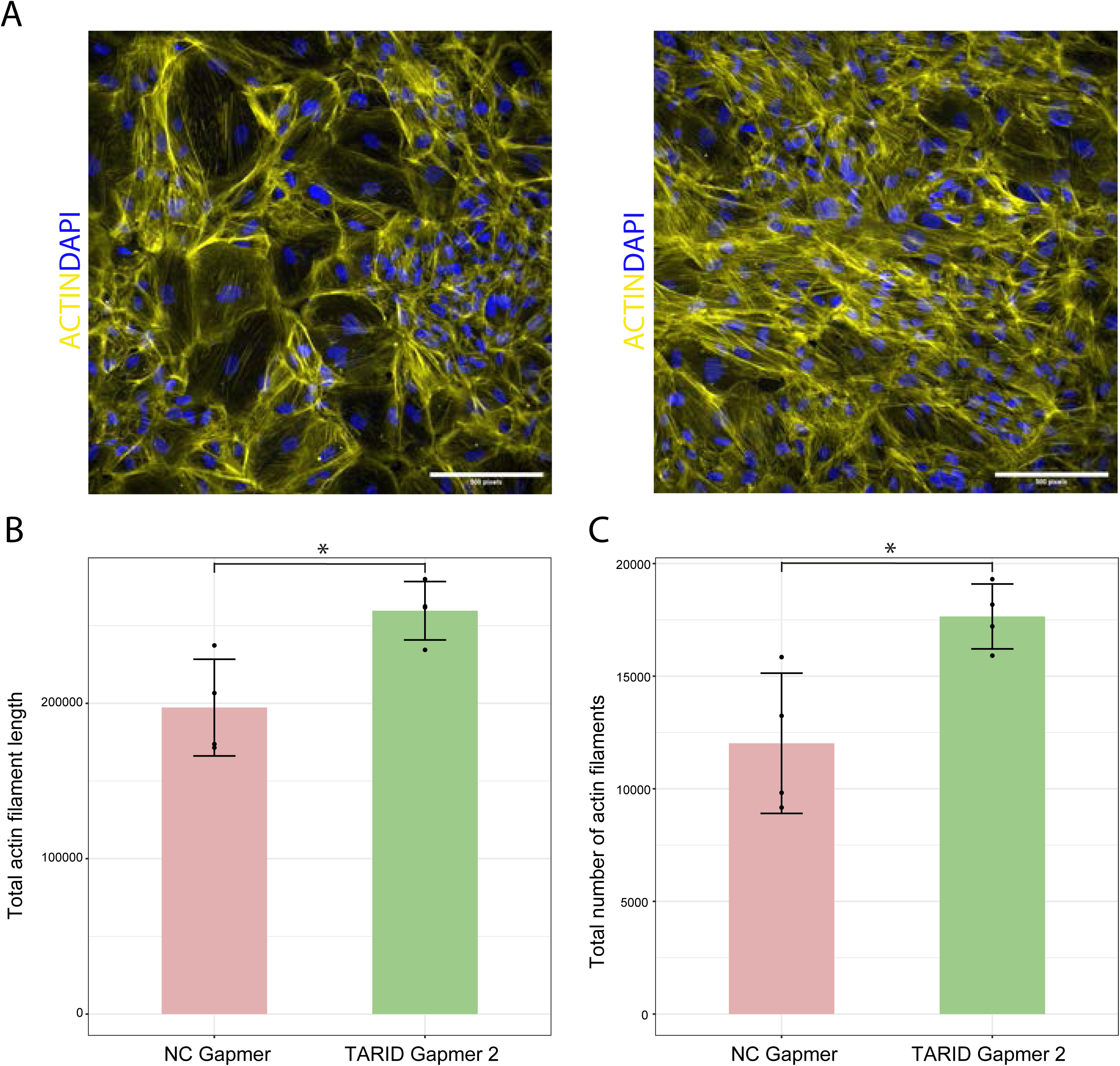
Actin staining analysis of podocytes following TARID knockdown. (**A**) Microscopic images of podocytes (CIHP-1) transfected with scrambled Gapmer (left) and TARID Gapmer (right) stained for actin (yellow) and DAPI (blue), bar = 500 pixels = 180 μm. (**B**) Quantification of total length of the actin filaments in podocytes (CIHP-1) treated with scrambled Gapmer (n=4) and TARID Gapmer (n=4). (**C**) Quantification of total number of actin filaments in podocytes (CIHP-1) treated with scrambled Gapmer (n=4) and TARID Gapmer (n=4).

## Discussion

The human transcriptome is predominantly composed of non-coding RNAs, with lncRNAs forming a significant subset [22]. LncRNAs are recognized for their varied roles in many biological processes such as cell differentiation and transcriptional regulation. However, the precise biological functions of the majority of lncRNAs are still largely unclear and a comprehensive understanding of lncRNAs in different kidney cell types in human kidney disease is lacking [4]. To explore this, we assessed our (publicly available) snRNA-seq dataset with a specific focus on lncRNAs in the human kidney [10], and demonstrated clear cell type and DKD specific expression of lncRNAs.

Most single-cell and single-nucleus RNA-seq studies have centered on the protein-coding genome, leaving a wealth of non-coding RNA biology largely unexplored. Interestingly, many lncRNAs exhibit tissue-specific expression patterns, often confined to a single cell type, which provides valuable insights into their specialized roles within cells [4]. It is believed that lncRNAs are often tissue-specific due to regulation by tissue-specific promoters and epigenetic marks, as well as specific signalling pathways active in certain tissues [23, 24]. They help regulate genes crucial for the identity and function of particular cell types [25]. Additionally, their proximity to tissue-specific genes and alternative splicing contribute to their specialized expression and roles [26]. This allows lncRNAs to participate in complex regulatory networks essential for cell specialization across different tissues [23].

A previous study that focused on lncRNAs in the mouse kidney found cell-type and age-specific expression of lncRNAs in the kidney [5]. However, there is a poor sequence conservation between human and mice for lncRNAs, which makes the understanding of the non-coding genome an exciting new area to understand human renal disease. As such, this study is the first to characterize lncRNAs across cell types in the human kidney, offering a valuable resource for understanding lncRNA expression patterns within this organ in humans. In total, we found that 10,327 lncRNAs are present in the snRNA-seq, which was 37% of the total amount of transcripts. We identified 349 lncRNAs with cell type-specific expression across all cell types in the kidney cortex, except for fibroblasts. These cell type-specific lncRNAs exhibited strong correlation with canonical protein coding RNA markers, suggesting that lncRNAs may serve as reliable biomarkers for distinguishing cell clusters. Notably, proximal tubular cells, which represent the largest cell population, displayed the highest number of cell type-specific lncRNAs. A significant positive correlation was observed between cell count and the number of cell type-specific lncRNAs across most cell types. Interestingly, podocytes presented an exception to this trend, with an unexpectedly high number of cell type-specific lncRNAs (42), indicating that these cells may depend on a unique lncRNA expression profile to support their specialized functions.

Furthermore, we identified 405 lncRNAs associated with DKD within a specific cell type, including 190 unique lncRNAs. We observed noticeable bias toward cells originating from female donors. Notably, *XIST*, an essential lncRNA responsible for X chromosome inactivation in females, was associated with DKD across multiple clusters. Since *XIST* is typically not expressed in males (except during germline development), this association may be biased by our study design, which compares three female controls to two females with DKD, potentially skewing results for lncRNAs located on the X chromosome [12]. Of the 405 lncRNAs associated with DKD, 104 exhibited cell type-specific expression. To demonstrate the relevance of our lncRNA resource, we further investigated the function of *TARID* as a case study.

Our focus on *TARID* stemmed from its unique association with the eGFR-related SNP, rs17643040-C, which is linked to reduced *TARID* expression in, among others, the kidney cortex and an increase in eGFR. *TARID* is notably upregulated in DKD and is specifically expressed in podocytes. Through network analysis, we identified 12 genes with strong correlations to *TARID*, including 8 that are crucial for podocyte function, highlighting its potential role in podocyte health and DKD. For instance, *PARD3B*, a key regulator of podocyte architecture, connects polarity signalling with actin regulation by modulating Rho-GTP levels, underscoring a functional pathway through which *TARID* may influence podocyte integrity [27]. Additional key genes associated with *TARID* include *ST6GALNAC3*, a recognized marker of podocyte injury [28], and *COL4A3*, mutations in which are linked to Alport syndrome. Mutations in *PLCE1* contribute to recessive nephrotic syndrome by influencing GTPase activity and actin dynamics in podocytes [29, 30]. Furthermore, *MAGI2* plays a vital role in preventing podocyte injury, progressive kidney disease, and renal failure [31]. These findings suggest that *TARID* and its network of associated genes are likely critical for maintaining podocyte function and kidney health, particularly in the context of DKD. Furthermore, pathway enrichment on genes correlated with *TARID* (Pearson correlation coefficient ≥ 0.4), showed enrichment for the *RHO GTPase cycle*. Additional pathway enrichment with KEGG, showed enrichment for *regulation of actin cytoskeleton*. Thus we hypothesized that *TARID* may be involved in the GTPase cycle to regulate actin cytoskeleton organization. This is a crucial component of podocytes as podocyte function is dependent on actin cytoskeleton regulation within the foot process [32].

This study is the first to establish a connection between *TARID* and podocyte function. *TARID* (*TCF21* antisense RNA inducing promoter demethylation) has been previously characterized in cancer, where it promotes activation of the tumor suppressor *TCF21*. In cancer cells, *TARID* recruits *GADD45A* to facilitate DNA demethylation, thereby reactivating *TCF21* expression in otherwise silenced cells [16]. *TARID* and *TCF21* are located adjacent to each other on the genome in opposite orientations. Our findings indicate that *TARID* knockdown slightly affects *TCF21* expression in podocytes. However, we found no correlation between *TARID* and *TCF21* expression in podocytes in the snRNA-seq.

We validated *TARID* expression in podocytes through reanalysis of two scRNA-seq datasets from untransplanted and transplanted kidney organoids and confirmed its localization with *in situ* hybridization in human kidney tissue sections [15]. Furthermore, knockdown of *TARID* in podocytes led to differential expression of 258 genes, among others we observed reduced expression of *WNT2* and *NOTCH3*. Both genes have previously been implicated in podocyte injury. Evidence suggests that the WNT pathway plays a critical role in podocyte injury, as WNT ligand expression is elevated in both murine experimental models for podocyte injury, as in human biopsy samples from individuals with glomerular diseases [33]. Similarly, *NOTCH3* is only expressed in injured podocytes, and *Notch3*-knockout mice, as well as wild-type mice treated with a *Notch3* inhibitor, showed protection against renal function decline [34]. *In vivo* studies have also linked *Notch3* activation in podocytes to morphological changes [35]. The overlap between genes correlated with *TARID* in the snRNA-seq data and the differentially expressed genes following *TARID* knockdown was minimal (3). This discrepancy may arise because correlation analyses and gene knockdown experiments measure distinct aspects of gene regulation, and the effects of knockdown may not always manifest directly. Nevertheless, we demonstrate that *TARID* knockdown significantly influences genes involved in diverse cellular processes, particularly actin cytoskeleton organization, aligning with the pathways associated with *TARID*-correlated genes identified in the snRNA-seq data. To further investigate this, we assessed the actin cytoskeleton following *TARID* knockdown and confirmed that *TARID* plays a critical role in actin cytoskeleton organization. We show that both the organization, length and total number of actin filaments is increased upon TARID knockdown. Based on these findings, we hypothesize that *TARID* knockdown may influence podocyte health by remodelling the actin cytoskeleton specifically in podocytes. It is noteworthy, however, that we prioritized *TARID* knockdown over overexpression, reflecting the expression patterns observed in our snRNA-seq results. Further studies are essential to confirm *TARID*’s role in podocyte function and to assess its potential as a therapeutic target.

### Conclusions

This study offers a valuable resource for examining lncRNA expression across various cell types in the human kidney. Through the identification of the podocyte-specific lncRNA *TARID*, we underscore the potential significance of cell type-specific lncRNAs in the pathogenesis of DKD.

## Author contribution

JAdK and RB designed the study and drafted the manuscript. JAdK conducted the analyses. JAdK, RJP, JHDP, HS, MK, LAKvdP, HJB and TG carried out the *in vitro* experiments, while RB, BDH, WCP, HW, and YM contributed to data acquisition and managed project logistics. All authors participated in data interpretation, critically reviewed the manuscript, and approved the final version. JAdK and RB serve as guarantors of the work.

## Data availability

Data sets have been deposited in the Gene Expression Omnibus and are available under accession number GSE195460, GSE151302 and GSE131882. The study also provides the processed count matrices for 11 snRNA-seq libraries, available under GEO GSE195460 [10]. All other data are included in the manuscript and/or Supplemental Data appendix.

