## Supplemental Material for "Cell Type-Specific and Diabetic Kidney Disease-Associated Expression of Long Non-Coding RNAs in Human Kidneys"

**Methods**

*Single-nucleus RNA sequencing*

We reanalysed our publicly available snRNA-seq dataset to focus on lncRNAs [1]. Eleven kidney cortex samples were obtained from control participants (n = 5) and patients with DKD (n = 6). Tissue samples were collected either post-nephrectomy for renal mass (n = 10) or from deceased organ donors (n = 1). We focused on lncRNAs annotated in GENCODE version 32 (GRCh38, Ensembl release 98), the same version used in the previous study, to capture the non-coding transcriptome. For consistency, we utilized the UMAP clustering from the original study [1]. Details on patient characteristics are also available in the original publication [1]. Cell-type-specific, defined by significantly higher expression in a single cell type compared to others, and DKD-associated (per cell type) lncRNAs, were identified using the Seurat *FindMarkers()* function, focusing on transcripts detected in at least 25% of cells. Statistical significance was determined using Bonferroni-adjusted p-values, with a threshold of FDR < 0.05. Gene regulatory networks were constructed by calculating Pearson correlation coefficients to identify associations between gene expression profiles. Pathway enrichment was conducted with Reactome Pathway Analysis (PA) and Kyoto Encyclopedia of Genes and Genomes (KEGG) using clusterProfiler (v4.12.0) and ReactomePA (v1.48.0). All snRNA-seq analyses were performed using R (v4.4.0) and the R package Seurat (v5.1.0).

*eQTL/GWAS eGFR signals*

To identify lncRNAs that may be causally related to kidney function, we used a publicly available Genome Wide Association Study (GWAS) (Catalog no. GCST90019506) of estimated glomerular filtration rate (eGFR). The GWAS included 349 significantly (P < 5 × 10^−8^) associated single nucleotide polymorphisms (SNPs) with eGFR. The Genotype-Tissue Expression (GTEx) database was used to screen for *cis* expression quantitative trait loci (cis-eQTLs).

*In situ hybridization*

Human Formalin-Fixed Paraffin-Embedded (FFPE) kidney tissue of 4-μm was stained with *in situ* hybridization (ISH). ISH was performed following the manufacturer's protocol (RNAscope™ Multiplex Fluorescent Reagent Kit v2). A custom probe for *TCF21* Antisense RNA Inducing Promoter Demethylation (*TARID)* on channel C3 was used (Hs-TARID-C3). To show podocyte specificity we used a probe for *WT1* on channel 2 (Hs-WT1-C2).

*Podocyte-specific expression of TARID in kidney organoids and other scRNA-seq datasets*

*TARID* expression was analysed in two independent scRNA-seq datasets. These datasets included untransplanted and vascularized transplanted kidney organoids from Koning et al [2] and are available in the ArrayExpress repository under accession number E-MTAB-11429. HiPSCs were differentiated into kidney organoids following a previously described protocol [2]. Briefly, hiPSCs were cultured as a monolayer for 7 days, then dissociated and transferred as 3D clumps onto a Transwell (0.4 µm pore polyester membranes). APEL2 medium with rhFGF-9 and heparin was added until 7+5 days. From day 7+5 onward, growth factors were removed, and the APEL2 medium was refreshed every other day for the remainder of the culture period. Organoids were cultured until transplantation in chicken embryos (day 7+12) or until day 7+20, coinciding with the harvest of transplanted organoids. ScRNA-seq data and metadata from kidney organoids—both untransplanted and transplanted—were processed using R (v4.2.1) with the Seurat R package (v4.2.0). Specifically, the data was grouped according to the “Final_Clusters” and “transplantation” metadata information. Scaled expression values and percentage of cells expressing *TARID* were obtained for each cell identity and transplantation condition using the *DotPlot()* function. The final dotplot was prepared with the ggplot2 package (v3.4.2) using the *geom_point()* function. Additional scRNA-seq datasets from the Kidney Interactive Transcriptome (KIT) database were examined [3]. These included publicly available datasets from human kidneys and kidney organoids.

*Additional glomeruli samples*

Micro dissected glomeruli from a frozen biopsy cohort of DKD patients, described by Baelde *et al*. (2007) were used to perform a quantitative real-time PCR (qPCR) to determine ***TARID*** levels [4]. These biopsies were obtained from the pathology archives of the LUMC and the Institute for Clinical Pathology, Heidelberg. All patients with DKD (n = 22) were diagnosed with type 2 diabetes. The control group (n = 10) consisted of the non-affected part of tumour nephrectomy samples or cadaver donor kidneys unsuitable for transplantation for technical reasons. The ***TARID*** RNA level were determined and corrected to the housekeeping genes glyceraldehyde-3-phosphate dehydrogenase (*GAPDH*), and hypoxanthine phosphoribosyl transferase (*HPRT*). The number of podocytes **was** quantified by calculating the mean of the gene expression levels of podocyte-specific genes *NPHS1*, *NPHS2*, and *WT1*. Sirius Red staining was used to quantify the amount of interstitial fibrosis and **served** as a histological marker for the progression of renal disease.

*In vitro experiments lncRNA TARID*

*TARID* expression was downregulated with three antisense Locked Nucleic Acid (LNA) oligonucleotides (Gapmers) targeting *TARID (*QIAGEN, custom design targeting *TARID)*. Human podocytes CIHP-1 (Ximbio) were cultured in 1640 Roswell Park Memorial Institute medium (RPMI, Gibco^TM^) containing 10% fetal bovine serum (FBS, Gibco^TM^), 1% insulin-Transferrin-Selenium (ITS-G, 100X, Gibco^TM^) and 1% Penicillin-Streptomycin at a temperature of 33 °C with 5% CO_2_. When cell density reached about 60%, CIHP-1 cells were exposed to Gapmers, Qiagen (three designs targeting *TARID, MALAT1* as positive control and a scrambled negative control) for 24 hours. Transfection of podocytes was performed using Lipofectamine LTX (Thermo Fisher scientific). After 24 hours the podocytes were thermos-switched to 37 °C with 5% CO_2_ for 14 days to fully differentiate.

*Quantitative real‑time polymerase chain reaction*

Quantitative real-time PCR (qPCR) was used to quantify the transcript expression of *TARID*, Transcription factor 21 (*TCF21)* and *MALAT1*. *GAPDH* was used as a housekeeping gene. The RNA from CIHP-1 cells was extracted with the RNeasy Micro Kit (Qiagen). Levels of RNA were quantified using a Nanodrop One Spectrophotometer (Thermo Fisher Scientific). Complementary DNA (cDNA) was synthesized with approximately 100 ng RNA using the Superscript RT II kit (Invitrogen). Individual gene expression levels were detected in triplicate using the SYBR Green master mix kit (Applied Biosystems). For this the following primers were used: *TARID*, forward (F) 5′‐GACTCACAGATCCAAGAATCCCA‐3′, reverse (R) 5′‐CAGCAGTTTGGCAAGATGGAG‐3′. *TCF21*, forward (F) 5′‐CATTCACCCGGTCAACCT‐3′, reverse (R) 5′‐TCAGGTCACTCTCGGGTTTC‐3′. MALAT1, forward (F) 5′‐GAATTGCGTCATTTAAAGCCTAGTT‐3′, reverse (R) 5′‐GTTTCATCCTACCACTCCCAATTAAT‐3′. *GAPDH*, forward (F) 5′‐GGAGCGAGATCCCTCCAAAAT‐3′, reverse (R) 5′‐GGCTGTTGTCATACTTCTCATGG‐3′.

*Bulk RNA sequencing*

RNA samples extracted from CIHP-1 podocytes treated with *TARID* Gapmer or control were sent for bulk RNA sequencing to Novogene. In short, messenger RNA (mRNA) was purified from total RNA using poly-T oligo-attached magnetic beads. After fragmentation, the first strand cDNA was synthesized using random hexamer primers followed by the second strand cDNA synthesis. The library was ready after end repair, A-tailing, adapter ligation, size selection, amplification, and purification. The library was checked with Qubit and real-time PCR for quantification and bioanalyzer for size distribution detection. Quantified libraries were pooled and sequenced on Illumina platforms, according to effective library concentration and data amount. Sequencing reads were aligned to the human genome (GRCh38 release 112) using STAR (v2.7.7a). Mapped reads were quantified for genomic features with featureCounts. Lowly expressed RNAs were filtered out (threshold of mean ≥5 copies) and read counts were normalized using the Trimmed Mean of M-values (TMM) method. For differential expression analysis, a quasi-likelihood negative binomial generalized log-linear model was applied using the edgeR package (v3.18) in R. Genes were considered differentially expressed if the contrast between *TARID* knockdown and control reached statistical significance, defined by a false discovery rate (FDR)-adjusted p-value of less than 0.05. Pathway enrichment was conducted with Gene Ontology (GO) using clusterProfiler (v4.12.0) and STRING [5]. All statistical analyses were performed in R (v4.4.0), and figures were generated with ggplot2 (v3.5.1) and enrichplot (v1.24.0).

*Immunofluorescent staining and imaging*

After full differentiation of podocytes and treatment with *TARID* and scrambled Gapmers, the cells were fixed with 4% paraformaldehyde and stained with Rhodamine Phalloidin (1:500; Invitrogen R415) to visualize F-actin and HOECHST 33258 (1:1000, Molecular Probes) for nuclear visualization. Blocking buffer contained 2% albumin from bovine serum (BSA) in PBS. Imaging was performed using a high-content confocal microscope (ImageXpress™ Micro Confocal, Molecular Devices) equipped with a 20× objective (Nikon Plan Apo Lambda; NA = 0.75). Detailed protocols for staining and imaging are described by Postma *et al* [6]. Images were quantified with ImageJ (v1.54m) Ridge Detection Plugin (v1.4.0) by measuring the total length and the number of the actin filaments. A Student's t-test was used to evaluate significance and a P-value < 0.05 was considered significant.

*Statistical analyses*

The correlation between *TARID* expression and Sirius Red staining was assessed using the Pearson correlation coefficient. Statistical significance of *TARID* upregulation in additional glomerular RNA samples and *TARID* knockdown in podocytes was evaluated using a Student's t-test. A P-value < 0.05 was considered significant.

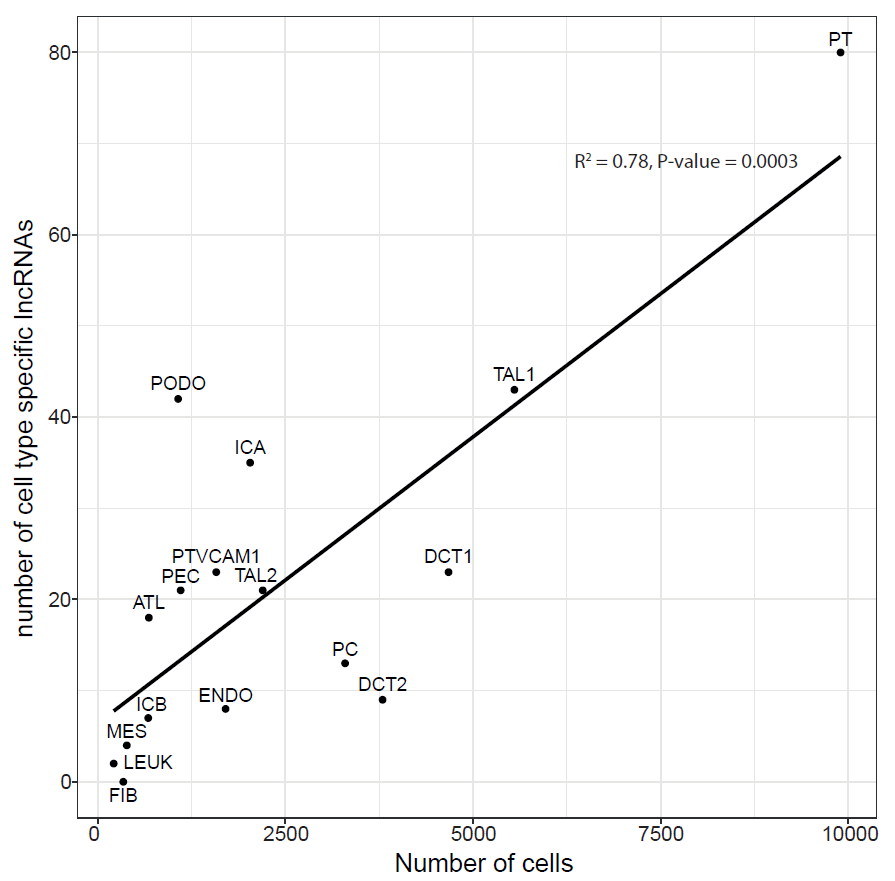

**ESM Figure 1. Correlation between number of cells and number of cell type specific lncRNAs in each cluster.** On the X-axis the number of cells in each cluster. On the Y-axis the number of cell type specific lncRNAs in each cluster. High correlation between number of cells and cell type specific lncRNAs in each cluster was observed, R^2^ = 0.78, P-value = 0.0003. However, some clusters, for example podocytes (PODO), do not follow this trend. In podocytes there are more lncRNAs cell type specific then expected based on the number of cell in this cluster.

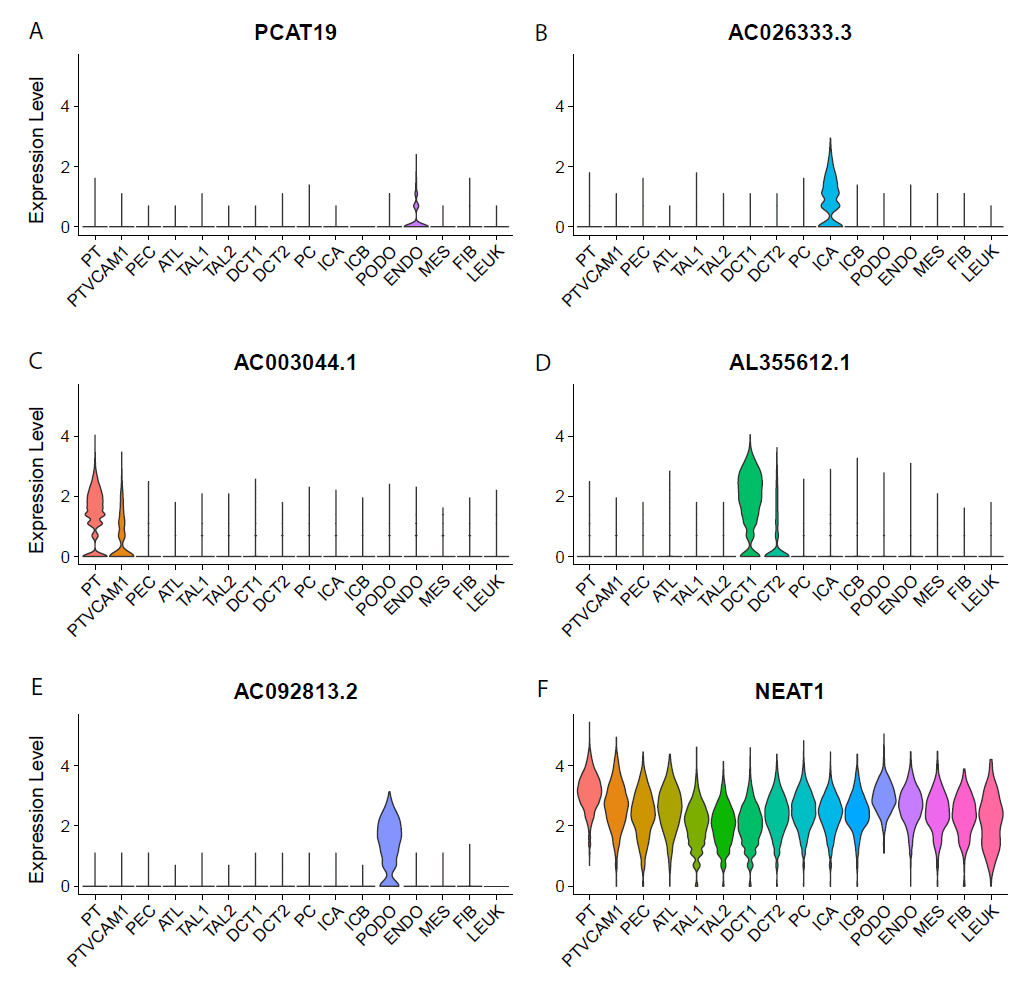

**ESM Figure 2. LncRNA expression in all cell types.** (**A**) lncRNA *PCAT19* is expressed in endothelial cells. (**B**) lncRNA *AC026333.3* is expressed in type A intercalated cells. (**C**) lncRNA *AC003044.1* is expressed in VCAM1-positive and -negative proximal tubule cells. (**D**) lncRNA *AL355612.1* is expressed in early and late distal convoluted tubule cells. (**E**) lncRNA *AC092813.2* is expressed in podocytes. (**F**) lncRNA *NEAT1* is expressed in all cell types but exhibits the highest expression in proximal tubule cells.

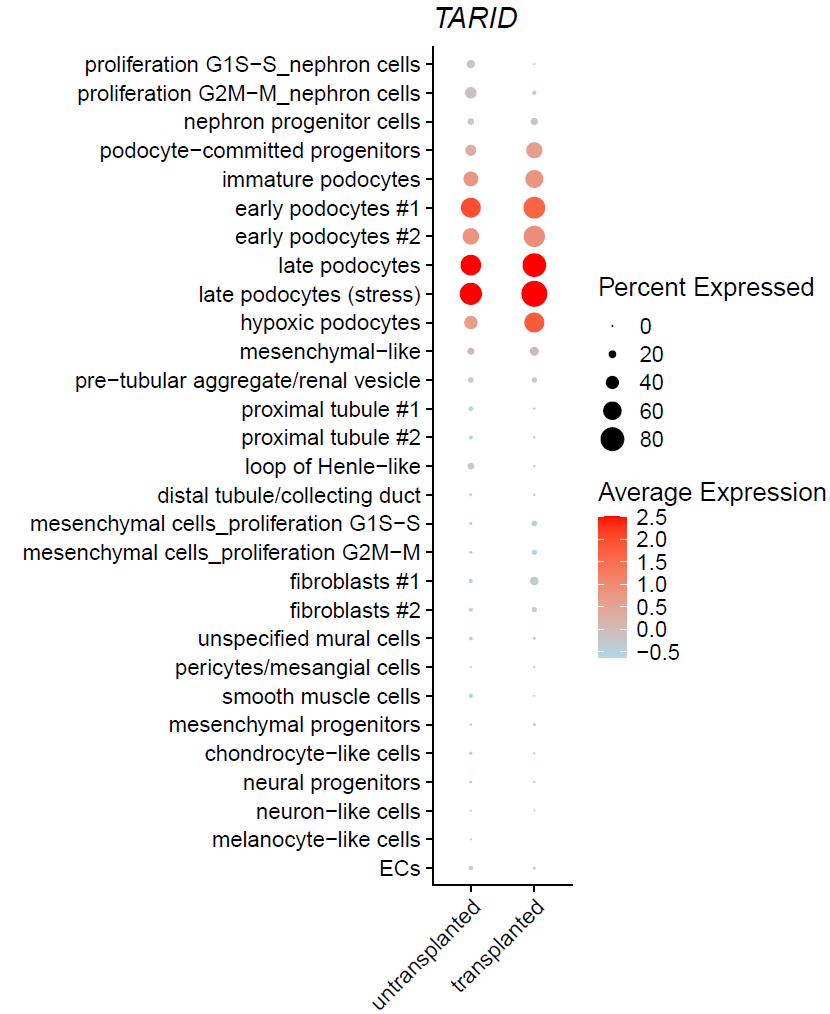

**ESM Figure 3. *TARID* is specifically expressed in podocytes of an untransplanted and transplanted kidney organoid.** LncRNA *TARID* is specifically expressed in two independent single cell RNA-seq datasets of an untransplanted and transplanted (for vascularization) kidney organoid. *TARID* is expressed in all developmental stages of podocytes (progenitor, immature, early, late, stressed and hypoxic).

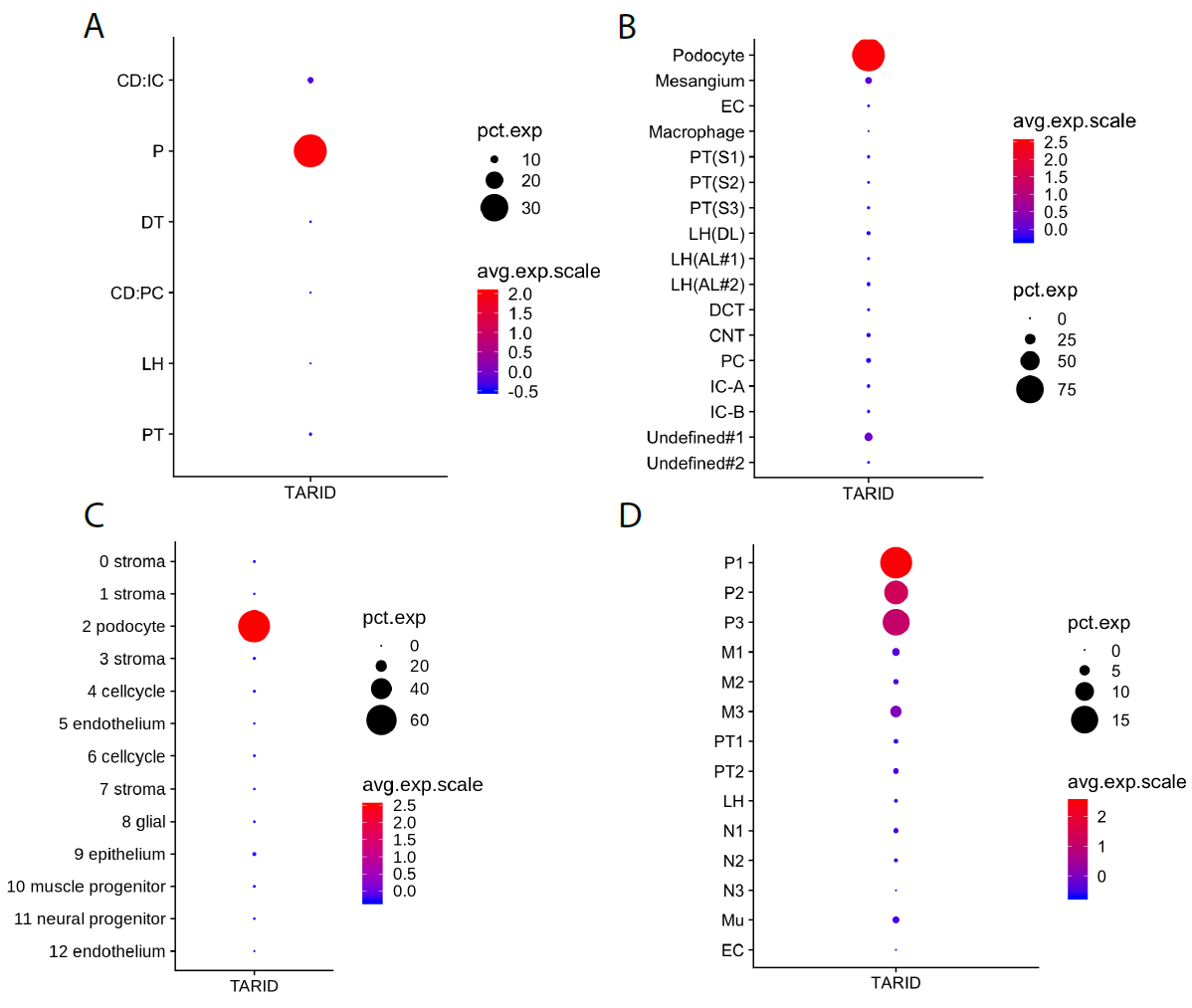

**ESM Figure 4. Podocyte-specific expression of *TARID* in four Kidney Interactive Transcriptomics (KIT) databases.** (**A**) *TARID* expression in KIT dataset: Healthy Adult Human Kidney - Epithelia: 4,297 nuclei. (**B**) *TARID* expression in KIT dataset: Healthy Human Adult Kidney - complete: 4,525 nuclei. (**C**) *TARID* expression in KIT dataset: Kidney organoids (Phipson et al 2019); 7,937 cells. (**D**) *TARID* expression in KIT dataset: Morizane's Kidney Organoid (d26): 15,191 cells.

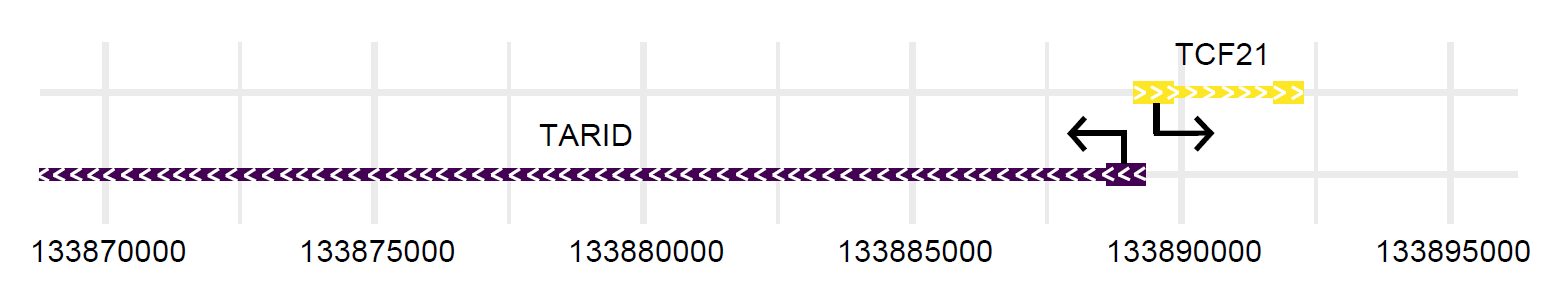

**ESM Figure 5. Genomic arrangement of *TARID* and *TCF21* on chromosome 6.** The figure shows the genomic positioning of *TARID* and *TCF21*, which are located adjacent to each other on chromosome 6 in opposite orientations.

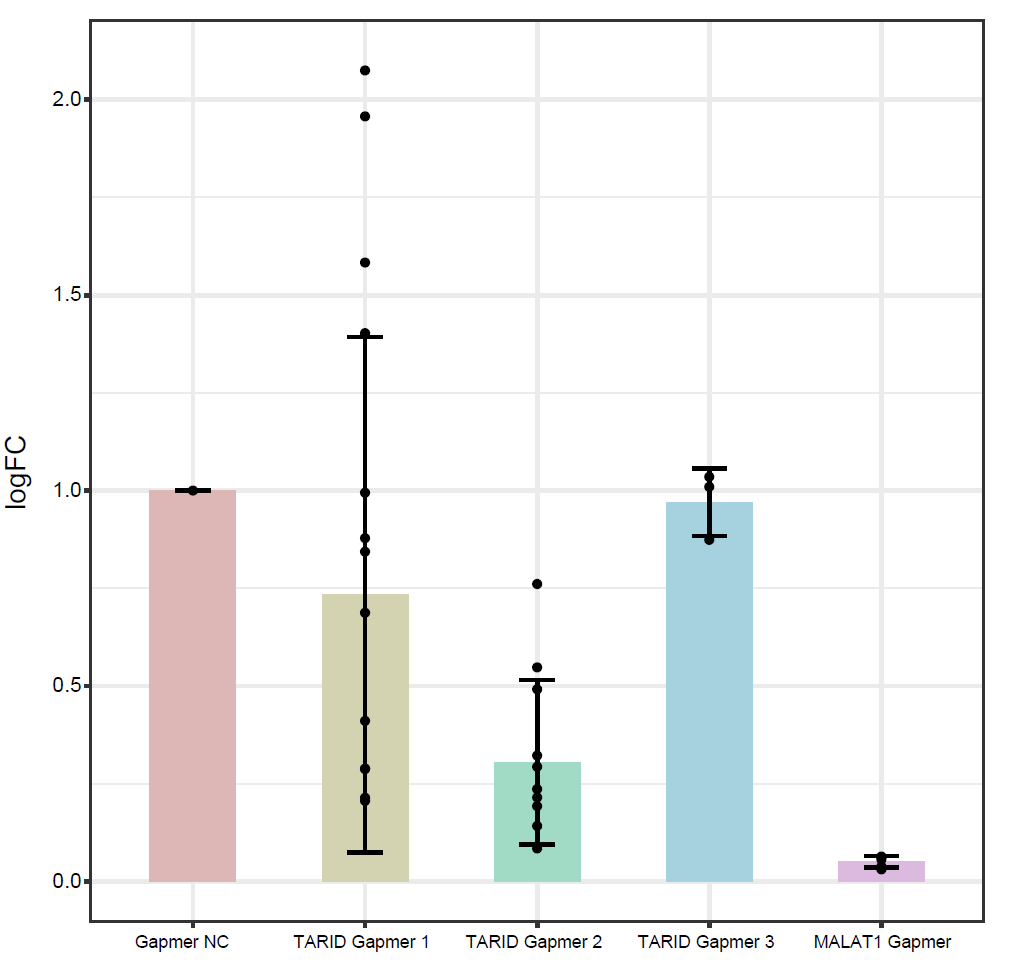

**ESM Figure 6. *TARID* knockdown in podocyte cell line with three different Gapmer designs.** Knockdown of *TARID* after introducing three different *TARID* antisense LNA Gapmers. Gapmer 2 seemed most effective in knockdown of *TARID*. Also, a positive control Gapmer *MALAT1* was introduced and showed effectively knockdown of *MALAT1* in podocytes.

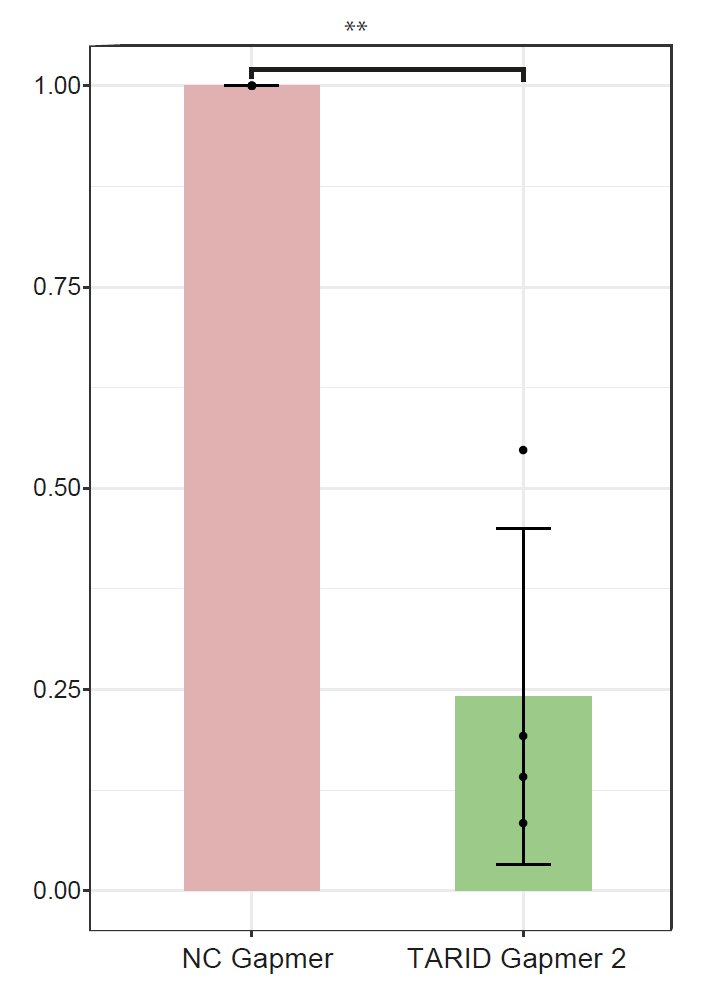

**ESM Figure 7. *TARID* knockdown in podocyte cell line with Gapmer 2.** Efficiently knockdown of TARID with Gapmer design 2 was observed. These eight samples were sent for bulk RNA-seq (NC Gapmer n=4, TARID Gapmer 2 n=4).

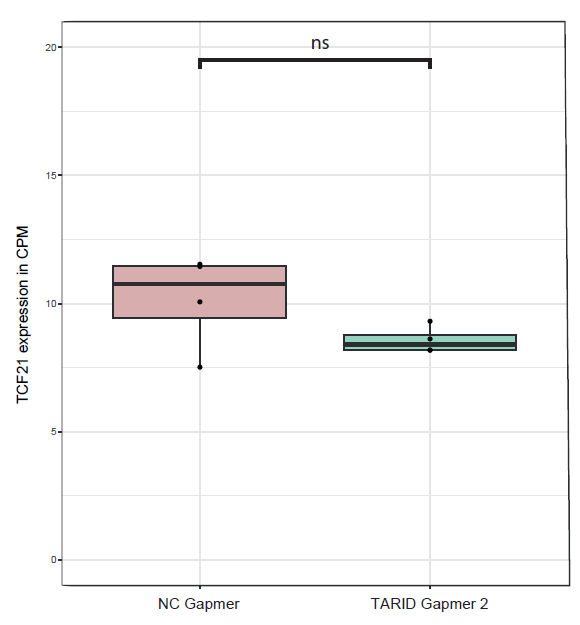

**ESM Figure 8. Bulk RNA-seq result of *TCF21* expression after *TARID* knockdown with Gapmer 2 in podocytes.** *TCF21* expression in the bulk RNA-seq of *TARID* knockdown in podocyte cell line.

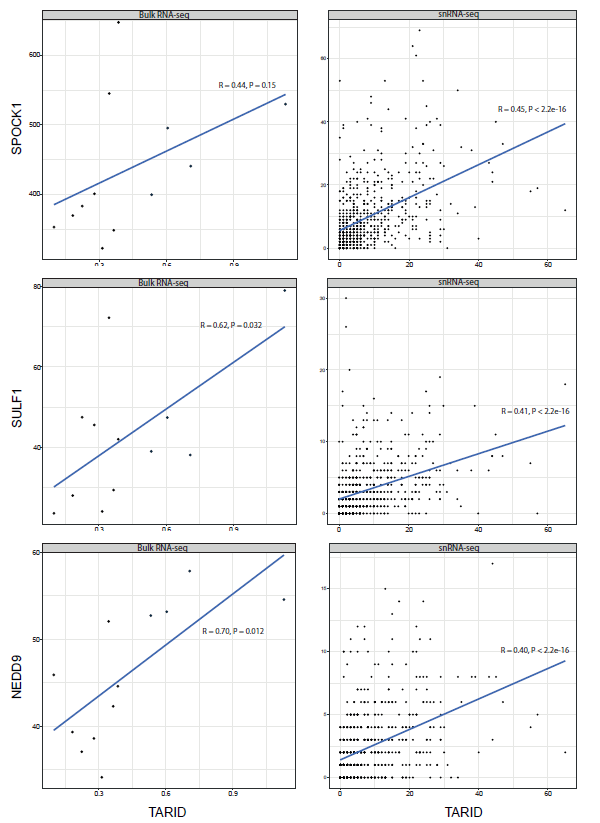

**ESM Figure 9. Correlation analysis of *TARID* with *SPOCK1*, *SULF1*, and *NEDD9* in podocyte RNA sequencing datasets.** The figure shows the correlation of *TARID* with *SPOCK1*, *SULF1*, and *NEDD9*, as observed in bulk RNA-seq of podocytes and in the podocyte cluster of snRNA-seq. Notably, three genes (*SPOCK1*, *SULF1*, *NEDD9*) exhibit significant association with *TARID* knockdown in podocytes and correlate with *TARID* expression in the snRNA-seq (Pearson correlation coefficient ≥ 0.4).

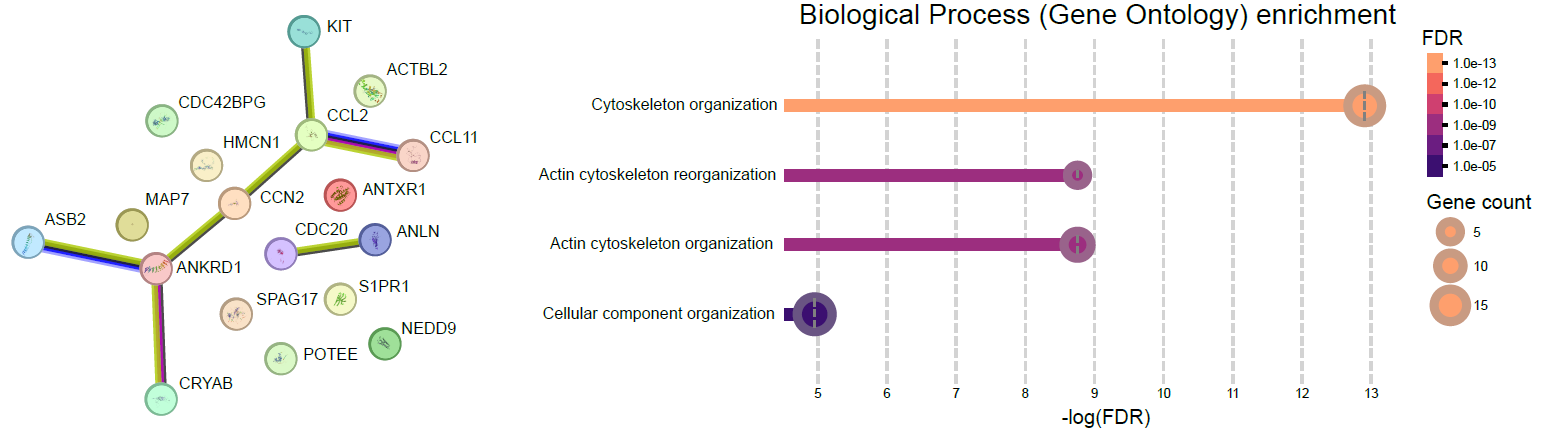

**ESM Figure 10. Investigation of the genes associated with cytoskeleton reorganization using STRING.** Figure shows pathway enrichment with STRING on the 19 genes associated with cytoskeleton organization from the 258 genes significantly associated with *TARID* knockdown in CIHP-1 cells.
